# Derepression masquerades as activation in a pentameric ligand-gated ion channel

**DOI:** 10.1101/2022.09.13.507804

**Authors:** Christian J.G. Tessier, Johnathon R. Emlaw, Raymond M. Sturgeon, Corrie J.B. daCosta

## Abstract

Agonists are ligands that bind to receptors and activate them. In the case of ligand-gated ion channels, such as the muscle-type nicotinic acetylcholine receptor, mechanisms of agonist activation have been studied for decades. Taking advantage of a reconstructed ancestral muscle-type β-subunit that forms spontaneously activating homopentamers, here we show that incorporation of human muscle-type α-subunits represses spontaneous activity, and furthermore that the presence of agonist relieves this α-subunit-dependent repression. Our results demonstrate that rather than provoking channel activation/opening, agonists may instead ‘inhibit the inhibition’ of intrinsic spontaneous activity. Thus, agonist activation may be the apparent manifestation of agonist-induced derepression. These results provide insight into intermediate states that precede channel opening and have implications for the interpretation of agonism in ligand-gated ion channels.

## Introduction

An enduring goal is to understand how agonists activate their receptors (Colquhoun, 2006a). For ligand-gated, or ‘agonist-activated’, ion channels, such as the nicotinic acetylcholine receptor (AChR), agonist binding culminates in activation/opening of an intrinsic ion-conducting pore (Colquhoun, 2006b). Early models postulated that agonists are effective at opening the channel because they have a higher affinity for the open state than the closed state, and thus their binding shifts the equilibrium towards the open state (Lape et al., 2008). In a landmark study, del Castillo and Katz proposed that agonism could be explained by a two-step process, where the agonist first binds to an inactive receptor, that then isomerizes to an active/open conformation (del Castillo and Katz, 1957). Within this framework, the ability of a given agonist to catalyze the isomerization step, and open the channel, was encoded in the chemical structure of the agonist, as well as its interactions with the receptor binding site. Full versus partial agonists differed in their ability to catalyze the isomerization step, with full agonists being more effective than partial agonists (Sine, 2012).

The advent of the patch clamp technique and the development of single-channel analysis methods have made detailed investigations of the activation mechanism possible (Colquhoun, 2007; Sine, 2012; Sivilotti and Colquhoun, 2016). Early single-channel experiments were consistent with del Castillo and Katz’s two step mechanism, however, to account for the two agonist sites on the AChR, the original scheme had to be extended to include two agonist-binding steps instead of one (Colquhoun and Sakmann, 1985). The emerging single-channel data also revealed additional complexity, including a class of brief closings, originally called ‘*nachschlag* shuttings’, which interrupted bursts of agonist-activated openings (Colquhoun and Sakmann, 1981). In some cases, the duration of these closings appeared to be agonist-independent, necessitating the introduction of an additional closed state in a more intricate scheme (Sine and Steinbach, 1986). These observations, combined with studies on the related glycine receptor (Burzomato et al., 2004) and continued technical improvements (Mukhtasimova et al., 2016), ultimately led to more elaborate mechanisms that incorporated intermediate closed states, termed ‘flipped’ or ‘primed’, which preceded channel opening (Lape et al., 2008; Mukhtasimova et al., 2009). These findings placed the origins of agonism at an earlier stage in the activation process (Lape et al., 2008), and demonstrated that the ultimate opening and closing rates of the AChR were independent of the agonist used to elicit them; a somewhat counterintuitive finding for a channel ‘activated’ by agonist.

Despite the explanatory power of intermediate closed states, they have been inferred solely from kinetic analysis of single-channel measurements, and thus they remain largely phenomenological (Colquhoun and Lape, 2012). What is ‘flipping’? What is ‘priming’? To gain further insight we have taken advantage of a reconstructed ancestral muscle-type AChR β-subunit, called ‘β_Anc_’, which at the amino acid level shares more than 70% sequence identity with the human β-subunit (Prinston et al., 2017). In addition to forming hybrid ancestral/human AChRs, where β_Anc_ replaces both the human β- and δ-subunits (Emlaw et al., 2021), β_Anc_ also forms homopentamers (Tessier et al., 2022). Despite presumably being devoid of agonist-binding sites, these β_Anc_ homopentamers ‘prime’ before opening spontaneously and, reminiscent of the wild-type AChR, display steady-state burst behaviour with *nachschlag-like* shuttings (Fig. 1A)(Tessier et al., 2022). For β_Anc_ to occupy both the β- and δ-subunit positions in hybrid AChRs, the (+) and (–) interfaces of β_Anc_ must not only be compatible with each other, but also with the corresponding interfaces of their bracketing α-subunits (Fig. 1A). This suggests that heteropentamers composed of β_Anc_ and the human α-subunit should be possible (Fig. 1B). If so, this would provide an opportunity to probe the function of the α-subunit, and specifically its role in agonism, in a unique context. Here we show that α/β_Anc_ heteromers are indeed viable, and that incorporation of the human α-subunit leads to an α-subunit dependent repression of spontaneous activity, which can be relieved by the addition of agonist. These findings provide insight into the origin of intermediate states that precede ligand-gated ion channel opening, and demonstrate how ‘agonist activation’ may instead be the apparent manifestation of agonist-induced derepression.

**Figure 1.**
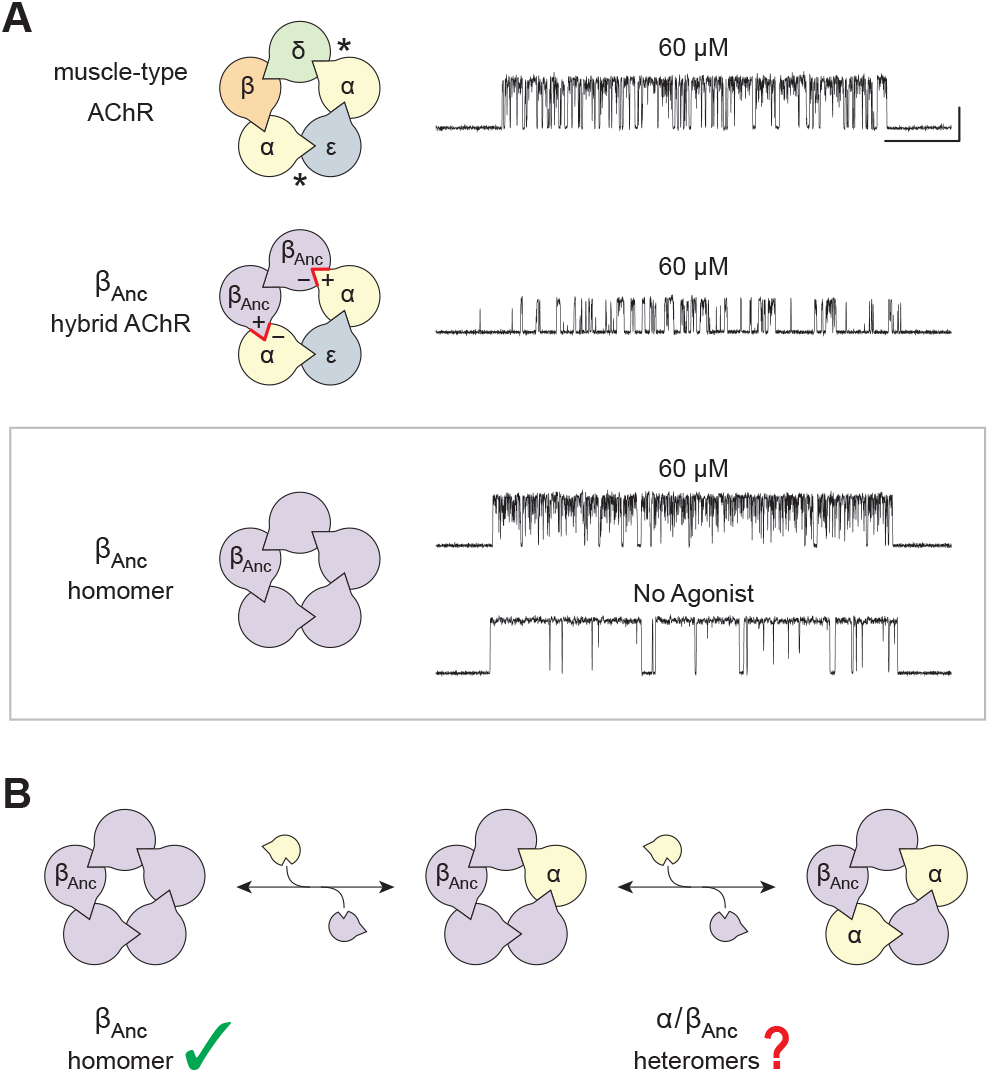
Subunit composition and single-channel activity of wild-type and β_Anc_-containing acetylcholine receptors. (**A**) Subunit stoichiometry and arrangement of the human adult muscle-type acetylcholine receptor (muscle-type AChR; *top*), where the agonist-binding sites at the α-δ and α-ε subunit interfaces are indicated with asterisks (*). A reconstructed ancestral β-subunit (β_Anc_; purple) forms hybrid acetylcholine receptors (β_Anc_ hybrid AChR; *middle*) where β_Anc_ substitutes for the human β-subunit (β; orange) and supplants the human δ-subunit (δ; green).For this to happen, the principal (+) and complementary (−) interfaces of β_Anc_ must be compatible with each other, as well as with the corresponding interfaces of their bracketing α-subunits (red highlight). β_Anc_ also forms homomers (*bottom;* boxed), which open spontaneously (no agonist), and whose single-channel activity in the presence of agonist mirrors that of the muscle-type acetylcholine receptor. Recordings were obtained in the cell-attached patch configuration with an applied potential of–120 mV and filtered with a 5 kHz digital Gaussian filter. Unless otherwise indicated, all recordings were acquired in the presence of 60 μM acetylcholine, where openings are upward deflections. The scale bar (20 ms, 10 pA) aligned to the top trace applies to all traces. (**B**) The compatibility of β_Anc_ with bracketing α-subunits in hybrid AChRs predicts that α/β_Anc_ heteromers are viable.

## Results

We previously showed that β_Anc_ forms spontaneously opening homopentamers that at the single-channel level share hallmarks of human muscle-type AChR activity, including steady state burst behaviour with *nachschlag-like* shuttings (Tessier et al., 2022). At concentrations of acetylcholine approaching saturation of the adult human AChR, bursts of single-channel activity from the two types of channels are essentially indistinguishable (Fig. 1A). This reflects both the high open probability of bursts of β_Anc_ homopentamer activity that is independent of agonist, as well as the similar extent of open-channel block by acetylcholine in the two types of channels (Tessier et al., 2022).

To determine if the human muscle-type α-subunit can coassemble with β_Anc_ to form α/β_Anc_ heteromers, we measured cell surface binding of radiolabeled α-bungarotoxin (α-Btx), an AChR competitive antagonist that can bind exclusively to determinants on the α-subunit (Dellisanti et al., 2007). When cells were cotransfected with cDNAs encoding the four human muscle-type AChR subunits, robust cell surface binding of α-Btx was detected (Fig. 2A). By contrast, cells transfected with cDNA encoding only the human α-subunit displayed little or no binding of α-Btx, indicating that on its own, the α-subunit is incapable of forming α-Btx binding sites that express on the cell surface. Similarly, when cells were transfected with cDNA encoding only β_Anc_, essentially no binding of α-Btx was detected. Given that single-channel activity of β_Anc_ homopentamers is readily observed (Fig. 2C, top trace)(Tessier et al., 2022), the simplest interpretation is that despite their robust cell surface expression, β_Anc_ homopentamers do not bind α-Btx. Evidently, as with its extant β-subunit counterparts, β_Anc_ lacks essential determinants of α-Btx binding. When cDNAs encoding β_Anc_ and the human α-subunit are transfected together, robust cell surface binding of α-Btx is restored, demonstrating that β_Anc_ can shepherd the α-subunits and their associated α-Btx binding sites to the cell surface, presumably by incorporating them into channels containing both types of subunits (Fig. 2A).

**Figure 2.**
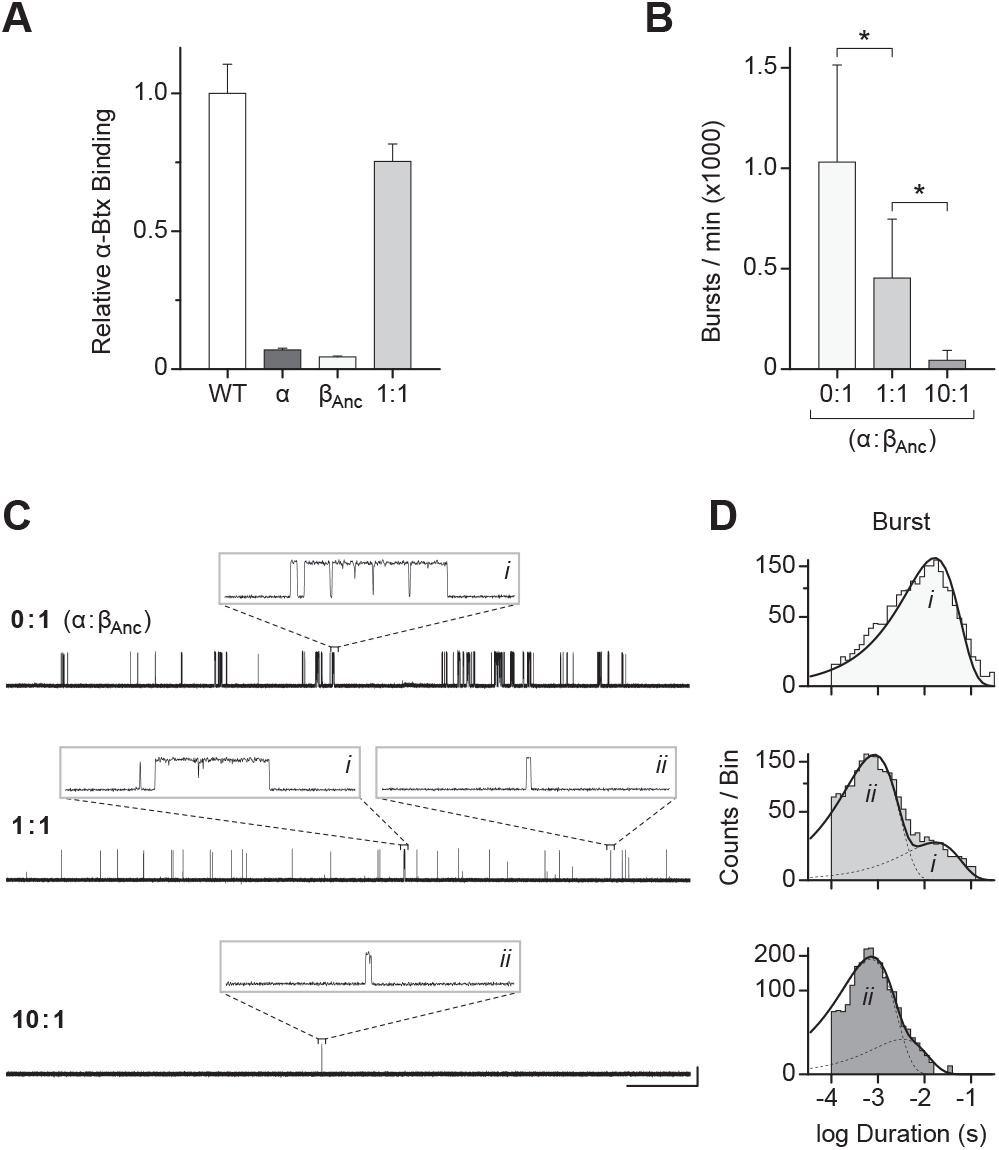
The human α-subunit and β_Anc_ can coassemble to form viable heteromeric channels that open spontaneously. (**A**) Relative cell surface binding of [^125^I]-α-bungarotoxin (α-Btx) to cells transfected with the full complement of cDNAs encoding the human adult muscle-type acetylcholine receptor subunits (WT; white), the α-subunit alone (α; darkest grey), β_Anc_ alone (β_Anc_; light grey), or a 1:1 (by weight) combination of the α-subunit and β_Anc_ (1:1; intermediate grey). Bar graphs represent the mean (error bars are plus one standard deviation) from two independent transfections, normalized to WT. (**B**) Frequency of spontaneous bursts of openings from cells transfected with cDNA encoding β_Anc_ alone (0:1; light grey), or with increasing amounts of human α-subunit cDNA (1:1, 10:1; by weight; intermediate and dark grey, respectively), while the amount of β_Anc_ cDNA is kept constant. Bar graphs represent the mean burst frequency (error bars are plus one standard deviation) averaged across ten separate single-channel patches, from at least three independent transfections. Difference between mean burst frequency is statistically significant (p < 0.05; asterisks), as determined by one-way ANOVA (Tukey’s multiple comparison test).(**C**) Single-channel burst behaviour of patches from cells transfected with different ratios (0:1, 1:1, 10:1; by weight) of α-subunit and β_Anc_ cDNA. In the absence of agonist, recordings were obtained in the cell-attached patch configuration with an applied potential of –120mV and filtered with a 5 kHz digital Gaussian filter. In each case openings are upward deflections, with the scale bar (2 s, 10 pA) beside the bottom trace applying to all zoomed out traces, and the inset boxes themselves representing 40 ms and 25 pA. (**D**) Burst duration histograms (see Methods) were manually fit (solid line) with a minimum sum of exponential components (dashed lines; labelled i and ii) containing the types of openings shown inset in (C).

To assess the functional consequences of replacing one or more β_Anc_-subunit(s) in β_Anc_ homopentamers with the human α-subunit, we cotransfected cells with cDNAs encoding both types of subunits and then examined the single-channel activity from cell-attached patches in the absence of agonist (Fig. 2B,C). Addition of α-subunit cDNA to a transfection mixture containing the same amount of β_Anc_ cDNA resulted in an overall reduction in the spontaneous activity of patches. The extent of this reduction related to the amount of α-subunit cDNA added and was best quantified as an α-subunit-dependent decrease in the frequency of spontaneous bursts (Fig. 2B).

Cotransfection with α-subunit cDNA also led to changes in the burst behaviour of the resulting channels (Fig. 2C,D). In the absence of the α-subunit, bursts were uniformly long, with a mean duration of ~10ms (Fig. 2C,D; top). When cells were cotransfected with α:β_Anc_ cDNAs at a ratio of 1:1 (by weight), the introduction of the α-subunit led to a second class of briefer events, bracketed by relatively long shut periods, which appeared as isolated openings (Fig. 2C; middle, inset *ii*) and contrasted with the bursts of closely spaced openings observed with β_Anc_ homopentamers (Fig. 2C; top & middle, inset i). The two types of events were clearly distinguished in burst duration histograms (Fig. 2D; middle), where individual openings separated by closings briefer than a specified critical closed duration (τ_crit_; 2 ms throughout) have been combined into bursts (Colquhoun and Hawkes, 1982). The heterogeneous burst behaviour of channels in these patches suggests that two types of channels, β_Anc_ homopentamers and α/β_Anc_ heteropentamers, are present. Consistent with this hypothesis, increasing the ratio of cotransfected α:β_Anc_ to 10:1 leads almost exclusively to isolated brief openings, essentially eliminating long bursts, and presumably β_Anc_ homopentamers. Evidently, cotransfecting the human α-subunit with β_Anc_ leads to a reduction in both the frequency and duration of spontaneous openings, and thus an overall repression of the spontaneous activity.

The above experiments show that the spontaneous single-channel activity of patches containing both β_Anc_ and the human α-subunit is reduced relative to those containing only β_Anc_ homopentamers, and thus that the presence of the α-subunits represses spontaneous activity. Given that the α-subunit only traffics to the cell surface in the presence of β_Anc_, a tacit assumption is that the reduced spontaneous activity is a result of the α-subunit replacing one or more β_Anc_ subunits in α/β_Anc_ heteromers. To determine how many α-subunits are present in spontaneously opening α/β_Anc_ heteromers we used an electrical fingerprinting strategy, where the α-subunit was tagged with mutations that reduce its contribution to overall single-channel amplitude. A similar strategy has been employed with tetrameric potassium channels (Niu and Magleby, 2002), and pentameric ligand-gated ion channels (Rayes et al., 2009; Andersen et al., 2011; daCosta and Sine, 2013; Andersen et al., 2013; daCosta et al., 2015), including the muscle-type AChR (Emlaw et al., 2021). Incorporation of one or more mutant low conductance α-subunits (α_LC_) would be expected to lead to a progressive decrease in the single-channel amplitude of resulting α_LC_/β_Anc_ heteromers, thereby allowing us to directly register the number of incorporated α_LC_-subunits in individual channels.

Using the wild-type AChR background, we confirmed that channels incorporating α_LC_ had a lower single-channel amplitude, but maintained a similar kinetic profile (Fig. S1). This control demonstrated that the conductance altering mutations did not affect other properties of the α-subunit. We then transfected cells with different ratios of α_LC_ and β_Anc_ cDNAs, and measured the distribution of single-channel amplitudes in patches from cells expressing the two types of subunits (Fig. 3). When cells were transfected with cDNAs encoding α_LC_ and β_Anc_, a variety of single channel amplitudes were observed in each patch (Fig. 3A,B). As expected, the relative proportion of high and low amplitude classes was dependent upon the ratio of transfected α_Anc_ to β_Anc_ cDNAs (Fig. S2). In mixtures of α_LC_ and β_Anc_ a maximum of only three amplitude classes were detected (Fig. 3C), where the mean amplitude of the highest amplitude class matched that of β_Anc_ homopentamers (Fig. 3C; inset 2, top). Thus, the simplest interpretation is that the intermediate and lowest amplitude classes originated from channels that have incorporated either one or two α_LC_ subunits, respectively. If channels containing more than two α_LC_-subunits were present in these patches, they did not open spontaneously, and thus were not evident in these fingerprinting experiments.

**Figure 3.**
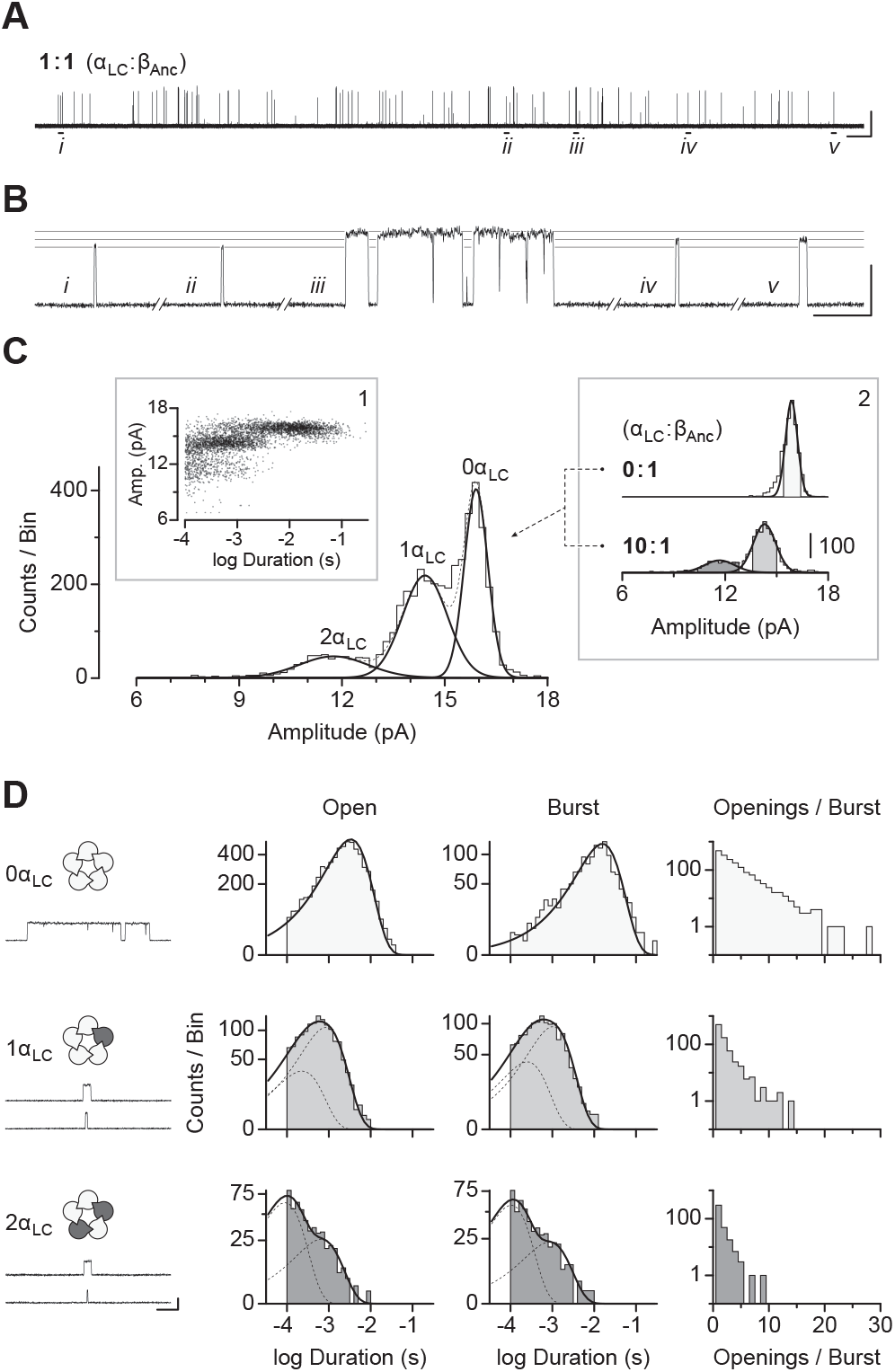
Electrical fingerprinting reveals the contribution of α-subunits to repression of spontaneous single-channel activity. (**A**) Co-transfection of a cDNA encoding the human α-subunit harbouring reporter mutations that reduce single-channel conductance (α_LC_) with cDNA encoding β_Anc_ leads to reduced spontaneous activity, where (**B**) the amplitude and duration of single-channel events is variable. Indicated openings in (A; *i-v*) are shown expanded in (B), with sight-lines overlaid to indicate the amplitudes of individual events. Openings are upward deflections, where the scale bars represent 1 s and 10 pA in (A), and 10 ms and 10 pA in (B). (**C**) Combining single-channel bursts from multiple recordings (see Fig. S2) where cells were transfected at different cDNA ratios (by weight) shows how single-channel amplitudes segregate into three well-defined amplitude classes (overlaid gaussians; solid lines), where each amplitude class corresponds to openings from channels incorporating either 0,1, or 2 α_LC_-subunits. Plotting the amplitude of individual bursts as a function of their duration reveals an apparent correlation (*inset 1*). (**D**) Dwell time analysis of selected bursts whose amplitude is within 1.5 standard deviations of the mean of each amplitude class (shaded regions from *inset 2* in (B)) reveals the contribution of each successive α_LC_-subunit (dark grey subunits in schematics) to open (*left*) and burst (*middle*) durations, as well as in determining the number of successive openings occurring in each burst (*right).* The traces under the schematics (left) show openings representative of each exponential component from visual fits of their corresponding duration histograms (*right*). Scale bar beside the bottom trace in (D) represents 5 ms and 10 pA. Recordings were obtained in the cell-attached patch configuration with an applied potential of –120 mV and filtered with a 5 kHz digital Gaussian filter.

To assess the impact of each α-subunit on the spontaneous activity of β_Anc_-containing channels we first plotted an amplitude versus duration scatter plot, where each data point represented the amplitude and duration of individual bursts in the fingerprinting experiments (Fig. 3C, inset 1). This analysis showed an apparent correlation between the amplitude and duration of bursts, with lower amplitude bursts being, on average, briefer in duration. This suggested that the incorporation of each successive α_LC_ subunit led to a progressive decrease in burst duration. To quantify this effect, we collected openings whose amplitude was within 1.5 standard deviations of the mean amplitude of each amplitude class (Fig. 3C, inset 2), and then performed dwell time analysis on each set of openings corresponding to spontaneous activity from channels with zero, one, or two α-subunits (Fig. 3D). Consistent with previous results (Tessier et al., 2022), spontaneous openings of β_Anc_ homopentamers were best fit by a single exponential component, and occurred in quick succession, ultimately coalescing into bursts that had a mean duration of ~10 ms. In contrast, channels containing one or two α-subunits exhibited two classes of spontaneous openings, both of which were briefer than the single class of openings in β_Anc_ homopentamers. When one α-subunit was present the longer of these two components was predominant, whereas when a second α-subunit was present the briefest component was predominant. Thus, with incorporation of each successive α-subunit the average duration of openings became briefer and briefer. In addition, when either one or two α-subunits were present, openings appeared as isolated events. This altered burst behaviour was quantified by binning bursts based upon the number of openings they contained, and showed a steep α-subunit-dependent drop off in the number of openings occurring within each burst (Fig. 3D; far right).

From the perspective of β_Anc_ homopentamers, the above data demonstrate that incorporation of human α-subunits leads to a repression of spontaneous single-channel activity. Given that in wild-type AChRs, residues from the α-subunits form the principal (+) face of the agonist-binding sites, we wondered whether this α-subunit-dependent repression could be relieved by the addition of agonist. We began by once again transfecting cells with various ratios of α:β_Anc_ cDNAs, and then recorded spontaneous single channel activity in the absence of agonist. To assess the impact of agonist, we identified cells that had a low, but quantifiable, spontaneous activity in the absence of agonist, and then returned to the same cell to record additional data from a new cell-attached patch, but this time in the presence of 300 μM acetylcholine. We reasoned that this high concentration of acetylcholine might be required to elicit a response given that α/β_Anc_ heteromers, with β_Anc_ providing the complementary (−) face of the agonist-binding site, lack important residues involved in agonist recognition (Emlaw et al., 2021). Furthermore, by returning to the same cell to acquire paired recordings, first in the absence, and then in the presence of acetylcholine, we were able limit patch-to-patch variability by controlling for the individual expression level of each transfected cell. The same strategy has been used to quantify calcium potentiation of the human α7 acetylcholine receptor (Natarajan et al., 2020). While extensive open-channel block resulting from the high agonist concentration precludes a meaningful comparison of dwell times (Fig. S3), we were able to assess the effect of agonist on burst frequency (Fig. 4). For β_Anc_ homopentamers (0:1; α:β_Anc_), although there was some variability in the frequency of bursts plus or minus agonist in paired patches from the same cell, the overall frequency of bursts was not significantly different in the presence or absence of agonist. By contrast, in cells transfected with a 10:1 ratio of α:β_Anc_ cDNAs (see also 1:1 in Fig. S4), paired recordings from the same cell revealed that, in each case, the presence of agonist led to a marked increase in the frequency of bursts. On average, the increase in burst frequency was more than 10-fold, revealing that agonist was able to relieve this aspect of the repression imparted by the α-subunits in α/β_Anc_ heteromers (Fig. 4C; Fig. S4).

**Figure 4.**
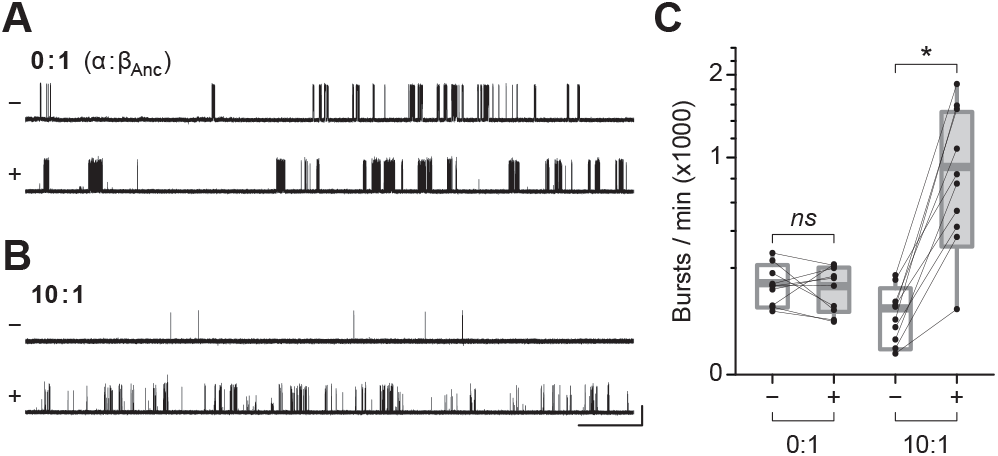
Agonist derepression of α/β_Anc_ heteromers. Single-channel activity from cells expressing (**A**) β_Anc_ homopentamers (0:1; by weight; α:β_Anc_) and (**B**) α/β_Anc_ heteromers (10:1) in the absence (−) and presence (+) of 300 μM acetylcholine. Recordings were obtained in the cell-attached patch configuration with an applied potential of −120 mV and Gaussian filter of 5 kHz. The scale bar beside the bottom trace in (B) applies to all traces and represents 1 s and 10 pA. (**C**) Comparison of burst frequency in paired recordings from cells expressing either β_Anc_ homopentamers (0:1), or α/β_Anc_ heteromers (10:1). In each case, paired cell-attached recordings from the same cell were acquired first in the absence (–), and then in the presence (+), of 300 μM acetylcholine in the patch pipette. Box plots represent one standard deviation from the mean, with the internal horizontal line denoting the mean of the 10 recordings in each case. Maximum and minimum values are presented as box plot whiskers.

## Discussion

Previously we showed that β_Anc_ readily forms homopentameric channels that open spontaneously (Tessier et al., 2022). Here we have shown that the human muscle-type α-subunit can coassemble with β_Anc_ to form α/β_Anc_ heteromers (Fig. 2A). Cotransfection of the α-subunit with β_Anc_ led to a decrease in the frequency of spontaneous openings in cell-attached patches (Fig. 2B), as well as alterations in the single-channel behaviour consistent with inhibition of the spontaneous activity (Fig. 2C). In both cases, the degree of inhibition was proportional to the amount of α-subunit cotransfected (Fig. 2). Tagging the α-subunits with conductance/reporter mutations allowed us to determine that α/β_Anc_ heteromers containing one or two α-subunits are viable, and open spontaneously (Fig. 3). Dwell time comparison of channels containing zero, one, or two α-subunits revealed that incorporation of each successive α-subunit progressively decreased the duration of both individual spontaneous openings, as well as bursts of closely spaced spontaneous openings (Fig. 3). Burst duration was further decreased through an α-subunit-dependent reduction in the mean number of openings occurring within each burst (Fig. 3). Finally, when both β_Anc_ and the α-subunit were cotransfected, the presence of agonist in cell-attached patches led to a dramatic increase in the single-channel activity (Fig. 4).

What do these experiments involving unnatural, heterologously expressed, spontaneously opening channels based upon a reconstructed ancestral muscle-type acetylcholine receptor β-subunit tell us about pentameric ligand-gated ion channels found in nature today? In particular, our experiments provide insight into the function of the α-subunits in muscle-type AChRs. Usually viewed as the principal subunits contributing to agonist binding and activation (Unwin et al., 2002), we’ve shown that the α-subunits also have the capacity to repress intrinsic spontaneous activity. Whether this is through actively repressing spontaneous openings, or by failing to contribute energetically to them in the same way that the β_Anc_ subunit they replace would, the result is the same: spontaneous openings are repressed in α/β_Anc_ heteromers. While this was demonstrated in an unnatural channel, the incorporated α-subunits were wildtype human α-subunits, raising the possibility that these α-subunits perform the same function in their natural setting, and repress intrinsic spontaneous activity in the heteromeric muscle-type AChR.

Mirroring muscle-type AChRs that contain two α-subunits, our fingerprinting experiments detected a maximum of two α-subunits in spontaneously opening α/β_Anc_ heteromers. While we cannot exclude the possibility that three (or more) α-subunits can exist in α/β_Anc_ heteromers, for this to happen, at least two α-subunits would have to occupy neighbouring positions in the resulting heteropentamer. This would require that the (+) and (–) interfaces of the α-subunit be compatible with each other, thereby allowing multiple α-subunits to self-associate. There is no evidence that the human muscle-type α-subunit can self-associate, and consistent with this, we did not to detect cellsurface binding of radiolabeled α-Btx when the α-subunit was transfected alone. The simplest interpretation is that α/β_Anc_ heteromers contain at most two α-subunits, where again mirroring muscle-type AChRs, the two α-subunits are separated by at least one intervening subunit (in this case a β_Anc_-subunit). Notably, the presence of two α-subunits in muscle-type AChRs is enough to ensure that spontaneous openings are infrequent (Nayak et al., 2012). Similarly, two α-subunits in α/β_Anc_ heteromers is enough to repress spontaneous activity, such that spontaneous openings occur infrequently in comparison to β_Anc_ homopentamers.

Modern mechanisms of AChR activation place the roots of agonism in ‘flipping’ or ‘priming’ steps that precede channel opening (Lape et al., 2008; Mukhtasimova et al., 2009, 2016). A consequence is that the ultimate opening and closing rates of the ‘agonist-activated’ AChR are, counterintuitively, independent of the agonist. The discovery that the α-subunits can repress spontaneous activity, and furthermore that this repression can be relieved by the presence of agonist, could reconcile these observations, and provide insight into phenomenological ‘flipping’ and ‘priming’ states. Instead of activating openings, agonists could instead derepress (i.e. ‘flip’ or ‘prime’) the channel, allowing it to open and close as dictated by its own intrinsic energy landscape. From this perspective, full versus partial agonists differ in their ability to derepress the channel, but once derepressed, the channel opens and closes in the same way irrespective of the efficacy of the bound agonist.

The possibility that the α-subunits hold the channel shut, as opposed to being responsible for opening it, also has potential implications for how agonism evolved in this family of proteins. Given that several aspects of wild-type AChR activation are independent of agonist and preserved in β_Anc_ homopentamers devoid of agonist-binding sites, it seems plausible that spontaneous channel activity existed before its regulation by agonist. In such a scenario, spontaneously opening channels evolved a subsequent layer of regulation through accumulating mutations that repressed their constitutive activity, which could then be restored by the presence of bound agonist. Essentially constitutive activity is inhibited by a repressor protein, in this case the α-subunits, which is then derepressed by the binding of an inducer, in this case an agonist (Fig. 5B). This is consistent with the observation that most AChR mutations alter the intrinsic tendency of the protein to open without changing the energetic contribution from bound agonist (Auerbach, 2012). If regulation by agonist evolved after spontaneous channel activity, constitutively active pentameric channels with biologically relevant functions should still be found in nature today, and indeed related GABA_A_ receptors containing the β3 subunit display spontaneous activity that is thought to control neuronal excitability (Sexton et al., 2021).

**Figure5.**
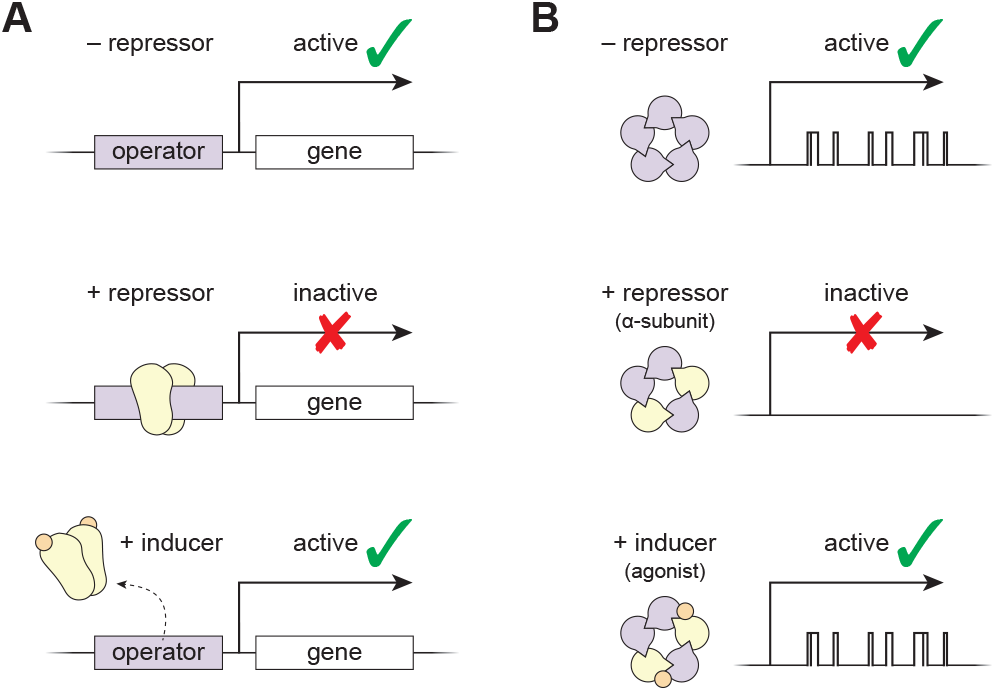
Parallels between induction of gene expression and agonism in a pentameric ligand-gated ion channel. (**A**) Constitutive expression of a gene (*top*) can be repressed by the binding of a repressor (yellow protein) to an operator sequence (*middle*), which can then be derepressed by the binding of an inducer (orange circle) to the repressor (*bottom).* (**B**) Constitutive activity of a pentameric ion channel (top; purple β_Anc_) can be repressed by incorporation of a repressor protein (*middle;* yellow α-subunits), which can then be derepressed by the binding of an inducer (bottom; orange agonist).

In famous work involving the *lac* operon, Jacques Monod and François Jacob showed that gene expression can be controlled by a repressor protein that binds to DNA and inhibits constitutive expression of a target gene (Fig. 5A). Expression of the target gene could then be induced by inhibiting the repressor protein. In Monod’s words, ‘the inducer acts not by provoking’ target gene expression, but instead ‘by inhibiting an inhibitor’ of its expression (Monod, 1966). Similarly, we have shown that rather than ‘provoking’ AChR openings, agonists may instead ‘inhibit the inhibition’ of constitutive activity, and thus ‘activation’ of the AChR, the prototypical ligand-gated ion channel (Changeux, 2020), could be the apparent manifestation of an analogous derepression mechanism (Fig. 5B).

## Methods

### Molecular biology

cDNA for the human AChR α1-subunit cloned into the pRBG4 plasmid was provided by Steven M. Sine, while cDNA encoding the ancestral β1 subunit (β_Anc_) was ordered as a synthetic oligonucleotide and cloned into the same pRBG4 plasmid (Prinston et al., 2017). Point mutations for the low conductance variant of the human α1 subunit (α_LC_; αD389R/N393R/A397R) were introduced into cDNAs by inverse PCR (Silva et al., 2017). Sanger sequencing confirmed the sequence of the entire reading frame.

### Mammalian cell expression

cDNAs encoding human and ancestral subunits were transfected into BOSC 23 cells (Pear et al., 1993). Cells were maintained in Dulbecco’s modified Eagle’s medium (DMEM) supplemented with 10% (vol/vol) fetal bovine serum at 37 °C, to a confluency of 50 to 70%. Calcium phosphate transfections were carried out for 3–4 h and terminated by media exchange. Experiments were performed 16–24 h after transfection. A separate plasmid encoding green fluorescent protein was included in all transfections to facilitate identification of transfected cells.

### Cell line authentication and mycoplasma testing

Approximately five million confluent cells were harvested and their total DNA isolated (E.Z.N.A.® Tissue DNA Kit), and then submitted to The Centre for Applied Genomics Genetic Analysis Facility (The Hospital for Sick Children, Toronto, Canada) for STR profiling using Promega’s GenePrint® 24 System. A similarity search on the 8,159 human cell lines with STR profiles in Cellosaurus release 42.0 was conducted on the resulting STR profile, which revealed that the cell line shares closest identity (88%, CLASTR 1.4.4 STR Similarity Search Tool score) with Anjou 65 (CVCL 3645). Anjou 65 is a child of CVCL 1926 (HEK293T/17) and is itself a parent line of CVCL X852 (Bartlett 96). Bartlett 96 is the parent line of BOSC 23 (Pear et al., 1993). PCR tests confirmed that the cells were free from detectable mycoplasma contamination (Uphoff and Drexler, 2011).

### Single-channel patch clamp recordings

Single-channel recordings were performed as previously described (Mukhtasimova et al., 2016). Briefly, recordings from BOSC 23 cells transiently transfected with cDNAs encoding wild-type, ancestral, or low conductance subunits, were obtained in the cell-attached configuration with a membrane potential of −120 mV and a temperature maintained between 19 and 22 °C. The bath solution contained (in mM) 142 KCl, 5.4 NaCl, 0.2 CaCl_2_ and 10 HEPES (4-(2-hydroxyethyl)-1-piperazineethanesulfonic acid), adjusted to pH 7.40 with KOH. The pipette solution contained (in mM) 80 KF, 20 KCl, 40 K•aspartate, 2 MgCl_2_, 1 EGTA (ethylene glycol-bis(β-aminoethyl *ether)-N,N,N’,N*-tetraacetic acid), and 10 HEPES, adjusted to a pH of 7.40 with KOH. Acetylcholine chloride (Sigma) was added to pipette solutions to the desired final concentration and stored at −80 °C. Patch pipettes were fabricated from type 7052 or 8250 nonfilamented glass (King Precision Glass) with inner and outer diameters of 1.15 and 1.65 mm, respectively, and coated with SYLGARD 184 (Dow Corning). Prior to recording, electrodes were heat polished to yield a resistance of 5 to 8 MΩ. Single-channel currents were recorded using an Axopatch 200B patch clamp amplifier (Molecular Devices), with a gain of 100 mV/pA and an internal Bessel filter at 100 kHz. Data were sampled at 1.0 μs intervals using a BNC-2090 A/D converter with a National Instruments PCI 6111e acquisition card and recorded by the program Acquire (Bruxton).

### Single-channel analysis

Single-channel event detection was performed using the program TAC (Bruxton), where data were analyzed with an applied 5 kHz digital Gaussian filter. Opening and closing transitions were detected using the 50% threshold crossing criterion, and corrected for instrument rise time using the R package *scbursts* (Drummond et al., 2019). Open and burst durations were placed into logarithmic bins (Sigworth and Sine, 1987), and the sum of exponentials was manually fit to the open and burst duration distributions within the program TACfit (Bruxton). Bursts of closely spaced openings were defined by a critical closed duration (τ_crit_), where individual openings separated by closings briefer than τ_crit_ (2 ms throughout) were concatenated with their intervening closings to produce individual bursts (Colquhoun and Hawkes, 1982). Duration histograms were constructed using a universal 2 ms burst resolution, and by omitting bursts briefer than 100 μs.

### Electrical fingerprinting

Cells were transfected as described above, but with an α_LC_ to β_Anc_ ratio of 10:1, 1:1, or 0:1 (α_Anc_:β_Anc_; by weight). Data were digitally filtered to 5 kHz within TAC and amplitudes were determined by setting the baseline and open-channel current for individual openings by eye. For each ratio, ~2000 events were collected and pooled from between 9-12 patches. A critical closed duration of 2 ms was uniformly applied throughout in order to define bursts of single-channel activity. To ensure that only fully resolved amplitudes were included in the analysis, events briefer than 100 μs were omitted. Pooled events were binned and plotted in event-based amplitude histograms, which were then fit with a set of Gaussian distributions. Data within 1.5 standard deviations of the mean amplitude encompassing each Gaussian, and representing each amplitude class, were then isolated for downstream dwell time analysis. Open and burst duration histograms were generated from events in each amplitude class in TACFit, and burst sorting and analysis was performed within R using the packages *scbursts* (Drummond et al., 2019) and *MASS* (Venables and Ripley, 2002).

### Paired recordings from the same cell

Paired recordings, in the absence (–) and then presence (+) of agonist, were obtained by successive cell-attached patches of the same cell. A 5 min continuous recording in the absence of agonist was obtained, and then a second 5 min recording from a new cell-attached patch, on the same cell, was acquired, but this time in the presence of 300 μM acetylcholine within the patch pipette. Data were filtered to 5 kHz, and all events within the 5 min window post voltage application were analyzed. A critical closed duration and a burst resolution of 2 ms, was uniformly applied throughout. Paired data plots were generated in R with the packages *ggplot2* (Wickham, 2016) and *ggpubr*.

### Radioligand binding experiments

AChR cell surface expression was measured by binding of [^125^I]-labeled α-bungarotoxin to transfected cells, as described previously (Emlaw et al., 2021). Briefly, approximately 850,000 BOSC 23 cells were plated onto 6 cm dishes and transfected with α, β_Anc_, or both α and β_Anc_ (1:1; by weight) cDNA’s. Cells were harvested one day post-transfection and incubated for 1 h at room temperature with [^125^I]-α-bungarotoxin (25 nm, specific activity of 10 Ci/mmol) in potassium Ringer’s solution. Ringer’s solution contained (in mM), 140 KCl, 5.4 NaCl, 1.8 CaCl_2_, 1.7 MgCl_2_, 25 HEPES, and 30mgL^-1^ bovine serum albumin, adjusted to pH 7.40 with KOH (daCosta et al., 2015). Cells were deposited onto 25 mm Whatman GF/C microfiber filter discs using a Hoefer filtration manifold and washed three times with 5 mL of Ringer’s solution. Nonspecific binding of [^125^I]-α-bungarotoxin to the filter discs was minimized by pre-incubating each disc in Ringer’s solution containing 1% bovine serum albumin for 1 h. Bound toxin was counted for 2 min in a Wizard2 1-Detector Gamma Counter (Perkin-Elmer).

## Supplementary Information

Supplementary Information is appended to this document, and includes Figures S1–S4.

## Acknowledgements

We thank Kathleen M. Gilmour for access to, and technical assistance with, a γ-counter. We thank Steven M. Sine, John E. Baenziger, and members of the dacosta]:[lab for comments on the manuscript. C.J.G.T. was funded in part by an Ontario Graduate Scholarship, while J.R.E. is the recipient of a Canada Graduate Scholarship from the Canadian Institutes of Health Research (CIHR) and a Natural Sciences and Engineering Research Council (NSERC) of Canada CREATE Scholarship. C.J.B.d.C. acknowledges grants from the Natural Sciences and Engineering Research Council of Canada (RGPIN-2016-04801), the Canada Foundation for Innovation (34475), the Canadian Institutes of Health Research (377068), as well as a New Frontiers in Research Fund-Exploration Grant (NFRFE-2018-00064). This manuscript was prepared using a modified version of a LATEX template kindly made available by Stephen Royle on GitHub (click here).

## Author Contributions

C.J.G.T. acquired and analyzed all electrophysiological data, while R.M.S. and J.R.E. acquired preliminary α/β_Anc_ heteromer recordings. J.R.E., C.J.B.d.C., and C.J.G.T. performed radiolabelled α-Btx experiments. C.J.G.T. and C.J.B.d.C. interpreted the data and wrote the manuscript. C.J.B.d.C. supervised the project.

## Author Declaration

The authors declare that they have no conflicts of interest with the contents of this article.

## Supplementary Information

**Figure S1.**
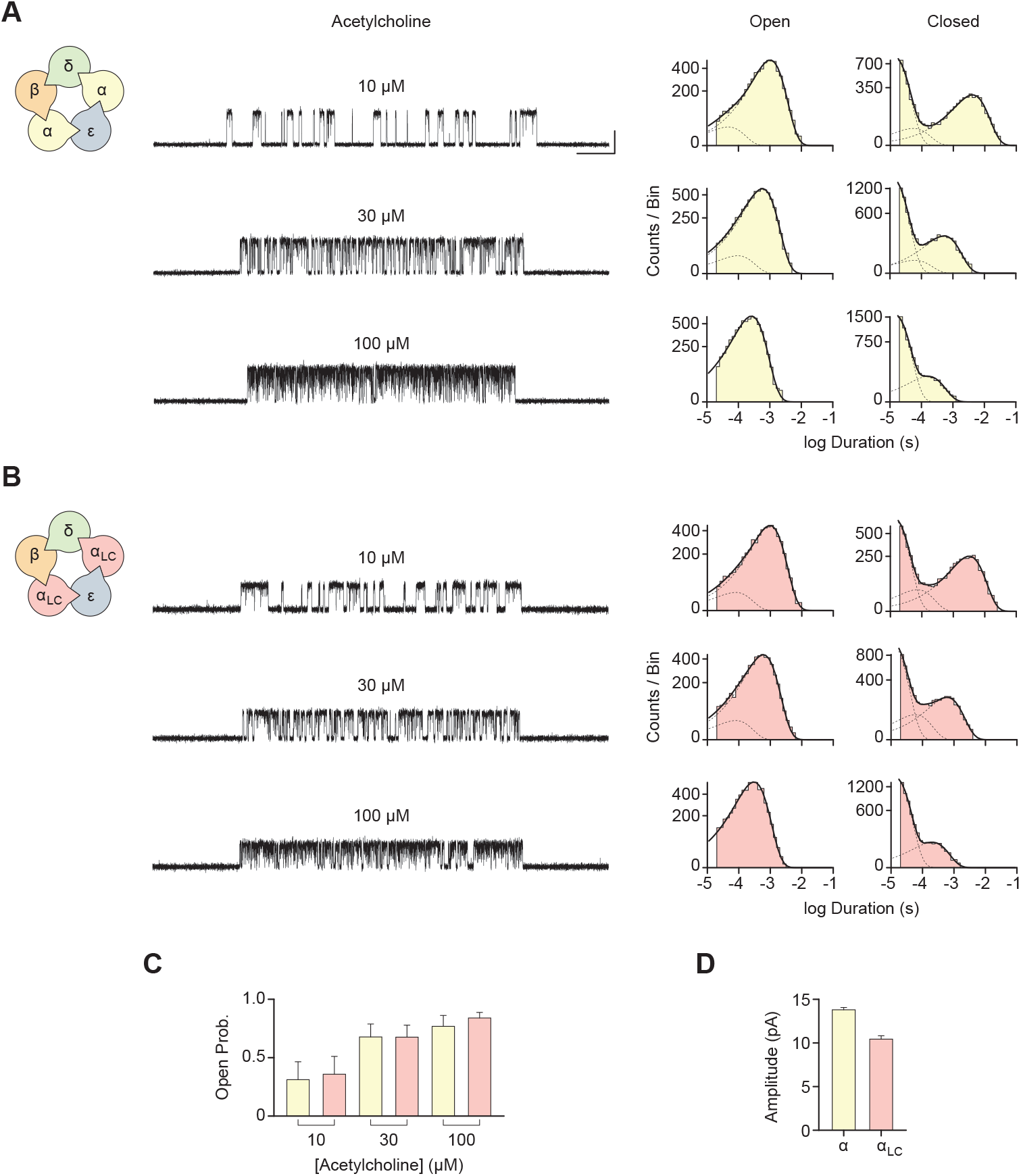
Human adult muscle-type acetylcholine receptors incorporating an α-subunit harbouring conductance mutations (α_LC_) have a lower single-channel amplitude, but maintain a similar kinetic profile. Subunit composition and single-channel burst activity of (**A**) wild-type and (**B**) α_LC_-containing human adult muscle-type acetylcholine receptors at the indicated acetylcholine concentrations. In each case, corresponding open and closed duration histograms are shown on the right, and fit (solid line) by a sum of individual exponential components (dashed lines). Recordings were acquired at −120 mV in the cell-attached patch configuration, and digitally filtered with a 10 kHz Gaussian filter. The scale bar in (A) represents 20 ms and 10 pA, and also applies to (B). (**C**) Mean open probability for wild-type (yellow) and α_LC_-containing (rose) channels is similar at the three acetylcholine concentrations. (**D**) Channels harbouring α_LC_ have a reduced single-channel amplitude. For (C), error bars represent one standard deviation from the mean, where each mean open probability was determined from a total of 75 bursts from three different patches (25 bursts each), from two separate transfections. For (D), error bars represent one standard deviation from the mean, where mean each amplitudes was determined from a total of 20 bursts from two different patches (10 bursts each), from two separate transfections.

**Figure S2.**
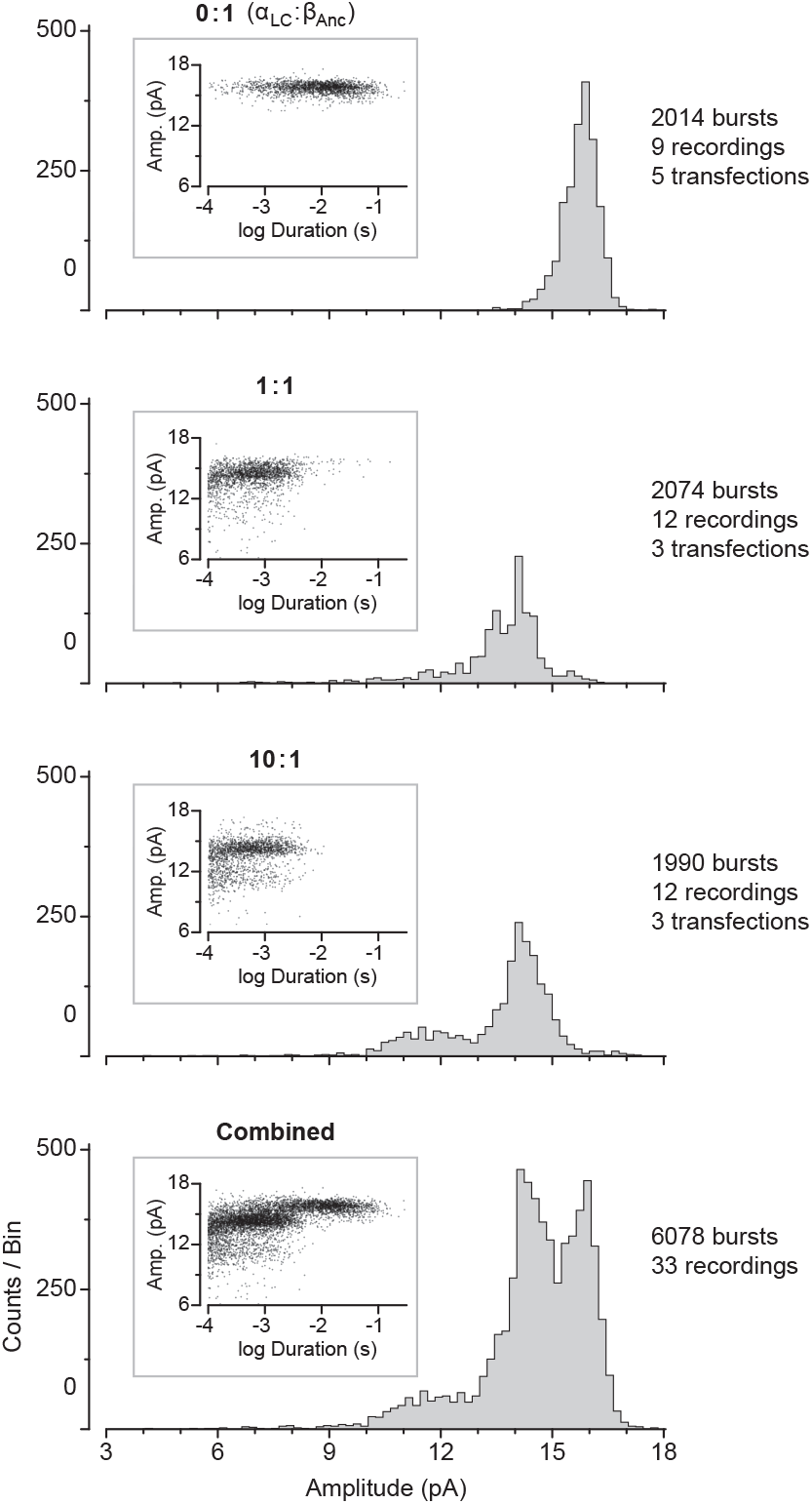
Amplitude distributions for single-channel bursts in patches where cells were transfected with the indicated ratios of α_LC_:β_Anc_ cDNA (by weight). As in Figure 3 of the main text, insets are plots of the amplitude of individual bursts as a function of their duration and reveal an apparent correlation. Combining bursts (*bottom*) from all three cDNA ratios (*top three plots*) reveal amplitude classes. As indicated, each distribution contains approximately 2000 bursts, from 9–12 individual recordings, from between 3–5 separate transfections.

**Figure S3.**
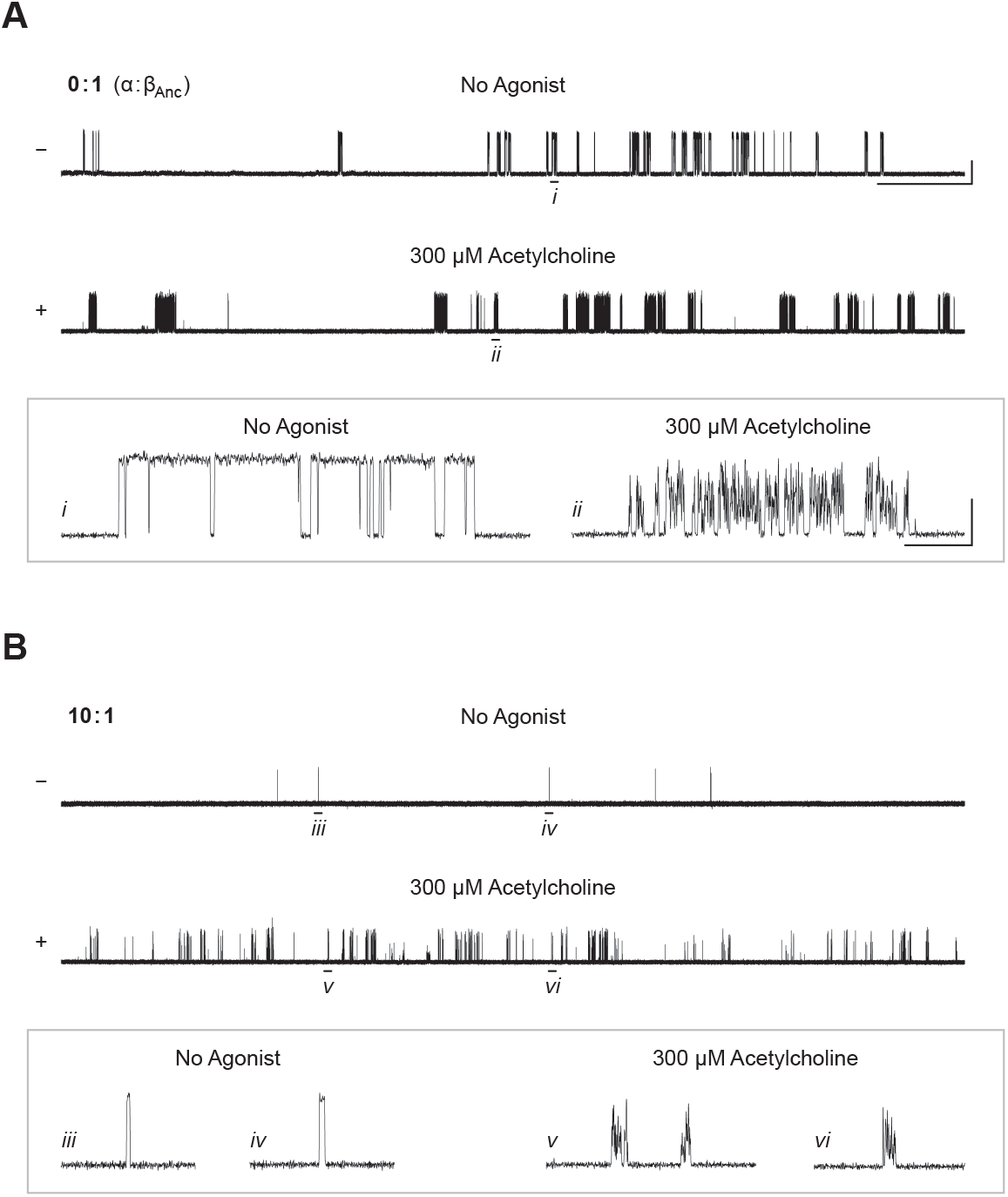
A high concentration of acetylcholine leads to extensive open-channel block. Single-channel activity from cells expressing (**A**) β_Anc_ homopentamers (0:1; by weight; α:β_Anc_) and (**B**) α/β_Anc_ heteromers (10:1) in the absence (−) and presence (+) of 300 μM acetylcholine. The top two traces in each panel are enlarged version of the traces presented in Figure 4 of the main text, but with select bursts highlighted (*i–vi*, below traces) and enlarged in the boxes below to show the effect of 300 μM acetylcholine. Recordings were obtained in the cell-attached patch configuration with an applied potential of −120 mV and Gaussian filter of 5 kHz. The scale bar beside the top trace in (A) applies to all zoomed out traces and represents 1 s and 10 pA, while the scale bar in the boxed region in (A) represents 10 ms and 10 pA and also applies to the boxed region in (B).

**Figure S4.**
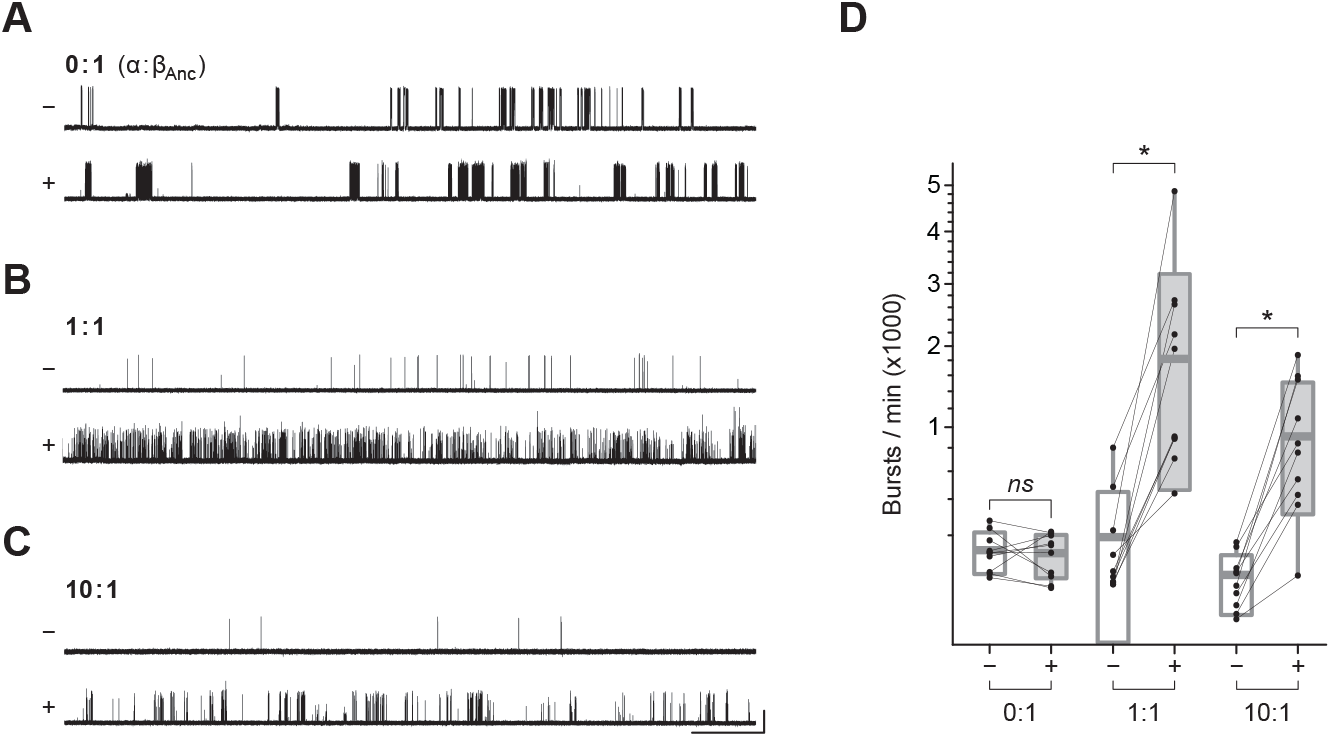
Agonist derepression of α/β_Anc_ heteromers (expanded). Figure 4 of the main text has been expanded to include cells transfected at an α:β_Anc_ cDNA ratio of (**B**) 1:1 (by weight). (**A**–**C**) The same cells were patched in the absence (–) and then in the presence (+) of 300 μM acetylcholine. Recordings were obtained in the cell-attached patch configuration with an applied potential of −120mV and Gaussian filter of 5 kHz. The scale bar beside the bottom trace in (C) applies to all traces and represents 1 s and 10 pA. (**D**) Comparison of burst frequency in paired recordings from cells expressing either β_Anc_ homopentamers (0:1), or α/β_Anc_ heteromers at two different α:β_Anc_ cDNA ratios (1:1 and 10:1). In each case, paired cell-attached recordings from the same cell were acquired first in the absence (−), and then in the presence (+), of 300 μM acetylcholine in the patch pipette. Box plots represent one standard deviation from the mean, with the internal horizontal line denoting the mean of the 10 recordings in each case. Maximum and minimum values are presented as box plot whiskers.

